# *Candida albicans* and *Staphylococcus aureus* reciprocally promote secretion of virulence factors

**DOI:** 10.1101/2024.09.25.615063

**Authors:** Raymond Pasman, Bastiaan P. Krom, Gertjan Kramer, Stanley Brul, Sebastian A.J. Zaat, Jianbo Zhang

## Abstract

Co-infections of *Candida albicans* and *Staphylococcus aureus* can significantly increase morbidity and mortality. This synergism is linked to the interactions between *C. albicans* and *S. aureus* that allow for staphylococcal co-invasion and dissemination. While it is known that extracellular virulence factors (ECVFs) contribute to this process, the effects of *C. albicans-S. aureus* co-culturing on ECVF composition remain unknown. In this study we used mass spectrometry-based proteomics to investigate the effect of co-culturing on the extracellular proteins released by the *S. aureus* and *C. albicans*. Co-culturing of *C. albicans* and *S. aureus* promoted the secretion of 7 cytolytic, 11 proteolytic, and 3 lipolytic ECVFs. Interestingly, co-culturing of *C. albicans* Als1p/Als3p mutant alleviated the increase for the majority of the differentially changed *C. albicans* ECVFs, but not for *S. aureus* ECVFs, highlighting the importance of Als1p/Als3p in the secretion of *C. albicans* ECVFs. Of 27 detected *S. aureus* ECVFs, 17 were significantly increased in co-culturing. Among these, maintenance of pH alone in *S. aureus* monoculture increased five haemolytic proteins, i.e., alpha haemolysin (Hly/Hla), beta haemolysin (Hlb), and gamma haemolysin (HlgA-C) to a similar extent as the co-culture. In contrast, maintenance of pH diminished the increase of protease-like proteins, (phospho)lipases, delta hemolysin, and leukotoxin, suggesting that both pH-dependent and pH-independent *C. albicans* factors affect *S. aureus* ECVFs. A cytotoxicity assay demonstrated that the secretome from co-culture has higher cytotoxicity towards human oral cells (Ca 9-22 and HO1N1) than monoculture. Finally, co-culturing increased the levels of non-extracellular virulence factors from both *C. albicans* and *S. aureus*. Taken together, the co-culturing of *C. albicans* and *S. aureus* reciprocally promotes their virulence potential, which may provide insights into the synergistic lethality during their co-infection *in vivo*.

## Introduction

*Candida albicans* is a commensal polymorphic fungus of the human oral, urogenital, gastro-intestinal, and skin mycobiome (1,2). In immunocompromised individuals, however, the fungus frequently becomes pathogenic (3). To become pathogenic, *C. albicans* need to switch from non-invasive yeast growth to invasive hyphal growth (4,5). Hyphal invasion comprises two key processes: the secretion of extracellular virulence factors (ECVFs) that damage the epithelium, allowing for easier invasion, and the physical invasion of hyphal cells, forcing the hyphae through epithelial layers (6–9). *C. albicans* hyphal invasion can result in a candidal bloodstream infection (BSI) (10). Interestingly, *C. albicans* BSIs frequently co-occur with bacterial bloodstream invasion (11), suggesting that the fungus facilitates co-invasion of bacteria. *Staphylococcus aureus* is the third most commonly isolated bacteria during *C. albicans* BSIs (11) and is the most common pathogen causing primary bacterial BSIs without an identified portal of entry or associated site of infection (12,13). Due to their co-isolation during candidal BSIs, *C. albicans* has been hypothesized to account for this lacking staphylococcal portal of entry.

Various *in vivo* murine studies have shown that *C. albicans* potently promotes co-invasion and dissemination of *S. aureus* and significantly increases lethality compared to monomicrobial infection (14–22). The *C. albicans* hyphal agglutinin sequence proteins 1 and 3 (Als1, Als3), the proteins responsible for *S. aureus* binding to hyphae, crucially contribute to *S. aureus* co-invasion (14–16).

Furthermore, the *C. albicans* ECVF candidalysin has been shown to significantly contribute to pathogenesis of *C. albicans/S. aureus* co-infections (16). Additional *in vitro* studies have proven that *C. albicans/S. aureus* co-culturing also increases the alpha toxin production of the *S. aureus* secretome by promoting the staphylococcal *agr* quorum sensing system in a pH dependent manner (23–26). Aside from candidalysin and staphylococcal alpha toxin, *C. albicans* and *S. aureus* secrete additional damaging ECVFs (cytolytic, proteolytic, and lipolytic) which could potentially contribute to invasion (**Table S1 and S2**). Furthermore, non-damaging ECVFs can also indirectly increase pathogenicity of both organisms during co-infection by aiding in immune evasion, adhesion, cell wall biosynthesis, and iron acquisition (**Table S1, S2**). However, the impact of co-culturing on these ECVFs remain to be elucidated. Therefore, the aim of this study was to identify which ECVFs are secreted by *C. albicans* and *S. aureus* and how co-culturing influence the virulence potential. We investigated the contribution of Als1/Als3 binding, biofilm integration, and *C. albicans* pH maintenance in mediating the changes in ECVF during co-culturing. Additionally, we tested whether co-culturing of *C. albicans* and *S. aureus* promoted ECVF cytotoxicity towards human oral squamous cells.

## Methods

### Strains and growth conditions

*C. albicans* and *S. aureus* strains are described in **Table 1**. In short, *C. albicans* and *S. aureus* strains were maintained on Sabouraud glucose agar supplemented with chloramphenicol (Sigma, 63567) and mannitol salt phenol red agar (MSA, Sigma, 89579), respectively. Single colonies were added to tryptic soy broth (TSB; Brunschwig Chemie, 211825) and cultured overnight at 37 ℃, 200 rpm. Cultures were rinsed with Dulbecco’s phosphate-buffered saline (DPBS; 137 mM NaCl, 2.7 mM KCl, 10 mM Na2HPO4, 1.8 mM KH2HPO4) and diluted to approximately ∼2 × 10^6^ CFU/ml for *C. albicans* and ∼2 × 10^7^ CFU/ml for *S. aureus* in Dulbecco’s Modified Eagle’s Medium (Sigma D5030) supplemented with 2.5 g/L dextrose, 1× Glutamax (Gibco 35050061), and 1× MEM non-essential amino acids (Gibco 11140050), with a final pH of 7.3 (mDMEM-DMP). Monocultures were generated by further diluting the culture in a 1:1 with mDMEM-DMP, while co-cultures were constituted by combining (undiluted) monocultures in a 1:1 ratio. For buffered growth mDMEM-DMP was supplemented with 100 mM of 4-(2-hydroxyethyl)-1-piperazineethanesulfonic acid (HEPES, Gibco 15630056).

**Table 1.** Strains of *C. albicans* and *S. aureus* used in this study

| Strain | Description | Reference |
| --- | --- | --- |
| *S. aureus* |  |  |
| ATCC12600 or NCTC8532 | wildtype | (27) |
| *C. albicans* |  |  |
| SC5314 | wildtype, clinical isolate | (28) |
| SC5314 *als1/als3* | *als1-1Δ::FRT/als1-2Δ::FRTals3-1Δ::FRT/als3-2Δ::FRT* | (29) |

### Biofilm growth and assessment

Mono- and co-culture biofilms were grown by inoculating wells of a 6 well plate (Corning 3506) with 3 mL of culture, prepared as described above, and incubating it stationarily for 2 hours at 37 ℃. Each well was washed for three times using DPBS to remove non-adherent cells. Finally, fresh mDMEM- DMP was added to the wells and all plates were incubated stationary for 72 hours at 37 ℃ in a humidified environment self-maintained with a flask of sterile water. Concerning trans well co-cultures, 1.5 mL of *C. albicans* mono culture was added to the inside of the 0.4 µm trans well inserts (Thermo Scientific, 140660) whereas 1.5 mL of *S. aureus* monoculture was added to the inside of the well plate. Trans well co-cultures were further treated identical to normal cultures with half the volume added on either side of the membrane. Medium pH was measured by pH 2.0-9.0 strips (Supelco, 1.09584) inoculated with 40 µL of medium. Culture medium was sampled for downstream analysis, as described below. The biofilms were collected using cell scrapers (Greiner, 391-3010), pelleted, and stored until further analysis using qPCR.

### Genomic DNA extraction and qPCR

Genomic DNA was extracted from the collected cell pellets using a DNeasy PowerBiofilm Kit (Qiagen, 24000) according to the manufacturer’s protocol. To determine the amount of fungal and bacterial DNA, a quantitative polymerase chain reaction (qPCR) was performed using primers specific to the fungal 28S rRNA gene (forward: GCATATCAATAAGCGGAGGAA AAG; reverse: TTAGCTTTAGATGATTTACCACC; probe: 6FAM-CGGCGAGTG AAGCGGSAARAGCTC-BHQ) (30) and bacterial 16S rRNA gene (forward: TCCTACGGGAGGCAGCAGT; reverse: GGACTACCAGGGTATCTAATCCTGTT; probe: 6FAM-CGTATTACCGCGGCTGCTGGCAC-BHQ1) (31). Fluorescence was measured using a LightCycler 480 System (Roche) and analysed using the corresponding system software.

### Protein concentration for proteomic analysis

To concentrate the secreted proteins in the spent media, the medium collected from six wells of the same culturing conditions were pooled and filtered using a 0.2-µm polyethersulfone (PES) filter (Sarstedt, 83.1826.001). The filtrate was further concentrated with 3 kDa Amicon ultra centrifugal filter (Pall, MAP003C37). The concentrated secretome samples were supplemented with protease inhibitors (Roche, 11873580001) according to manufacture manual for proteomic analysis or immediately stored without protease inhibitor supplementation for cytotoxicity assay. All secretome samples were stores at -80 ℃ until use.

### Sample preparation for LC-MS analysis

Samples were prepared and measured according to Šimkovicová *et al*. (2024) (32). In short, samples were thawed, reduced, and alkylated by incubation with tris-(2-carboxyethyl)phosphine (10 mM) and chloroacetamide (30 mM) (Sigma Aldrich) for 30 minutes at 70°C. Next, samples were prepared for mass spectrometry analysis using the single-pot, solid-phase-enhanced sample preparation (SP3) protocol (33). Soluble protein recovery was optimized by ensuring no detergents were added to the samples and precipitation time was extended to 30 minutes (room temperature). Beads used for washing were air-dried and resuspended in ammonium bicarbonate (100 mM, Sigma Aldrich) after which trypsin (Sequencing Grade Modified, Promega) was added at a protease-to-protein ratio of 1:50 (w/w) at 37°C. Formic acid was added to the overnight digestion at a final concentration of 1% and pH of 2. Finally, peptides were recovered using a magnetic separator device.

### LC-MS analysis for quantitative proteomics

Samples were separated by reversed phase chromatography using an Ultimate 3000 RSLCnano UHPLC system (Thermo Scientific, Germeringen, Germany) and peptides were separated using a Aurora Ultimate 75 μm × 250 mm C18 column (, 1.6 μm particle size, Ionopticks, Australia), maintained at 50 °C and operated at a flow rate of 400 nL/min with 3% solvent B for 3 minutes (solvent A: 0.1% formic acid in water, solvent B: 0.1% formic acid in acetonitrile, ULCMS-grade, Biosolve). Next, a multi-stage gradient was applied (17% solvent B at 21 minutes, 25% solvent B at 29 minutes, 34% solvent B at 32 minutes, 99% solvent B at 33 minutes, kept at 99% solvent B till 40 minutes). For equilibration, the system was returned to initial conditions (t = 40.1 minutes) for 58 minutes. Eluted peptides were electrosprayed by a captive spray source via the column-associated emitter and were analyzed by a TIMS-TOF Pro mass spectrometer (Bruker, Bremen, Germany) operated in PASEF mode for standard proteomics acquisition. MS/MS scans were initiated 10× with a total cycle time of 1.16 seconds, a target intensity of 2•10^4^ , an intensity threshold of 2.5•10^3^, and a charge state range of 0-5. Active exclusion was enabled for a period of 0.4 minutes and precursors reevaluated when the ratio of current intensity: previous intensity exceeded 4.

### Spectral data processing and proteome database search

LC-MS data were processed using MaxQuant software (version 1.16.14.0) using standard settings, i.e. trypsin/p as the enzyme allowing for 2 missed cleavages with carbamidomethylation at cysteine as a fixed modification and oxidation at methionine as a variable modification searching the proteome databases of: Candida_albicans_UP000000559 and Saureus_UP000008816. MaxQuant outputs were used for subsequent analysis using Perseus (version 2.0.7.0). Proteins that are only identified by peptides carrying one or more modified amino acids as well as reverse and potential contaminant proteins were removed from the dataset. The remaining data was log2 transformed and the proteins that were not detected in both samples of at least one condition were removed. Next, the remaining proteins were annotated using the 2019_11 release of the Max Planck institute of biochemistry annotation database (*C. albicans* SC5314 and *S. aureus* NCTC 8325) after which the dataset was split into two sets; one set containing all *C. albicans* proteins and one set containing all *S. aureus* proteins. For both sets the missing values were imputed based on the low end of the corresponding normal distribution (width = 0.3, down shift = 1.8) and all values were subtracted with the most frequent value. Protein differences were tested on significant using ANOVA with a permutation-based FDR of 0.05 and 250 number of randomizations after which non-significant proteins were removed. Principal component analysis was used to identify sample group differences based on the remaining proteins. Finally, the remaining data was normalized using Z-score normalization and significantly differing proteins between monoculture and co-culture conditions were visualized using volcano plots based on a Pearson correlation with an FDR of 0.05. Using the virulence factor database (34), Aureo Wiki (35), Uniprot (36), STRING database (37), and various studies (38–40), extracellular (ECVFs) and non-extracellular (N-ECVFs) virulence factors were identified. Results were visualized using Microsoft Excel (version 2402, build 17328.20282).

### Data analysis

Using Perseus (v2.0.7.0) the proteins that are only identified by peptides carrying one or more modified amino acids as well as reverse and potential contaminant proteins were removed from the dataset. The remaining data was log2 transformed and the proteins that were not detected in both samples of at least one condition were removed. Next, the remaining proteins were annotated using the 2019_11 release of the Max Planck Institute of Biochemistry annotation database (*C. albicans* SC5314 and *S. aureus* NCTC 8325), after which the dataset was split into two sets; one set containing all *C. albicans* proteins and one set containing all *S. aureus* proteins. For both sets missing values were imputed based on the low end of the corresponding normal distribution (width = 0.3, down shift = 1.8) and all values were subtracted with the most frequent value. Protein differences were tested on significant using ANOVA with a permutation-based FDR of 0.05 and 250 number of randomizations after which non-significant proteins were removed. Principal component analysis was used to identify sample group differences based on the remaining proteins. Finally, the remaining data was normalized using Z-score normalization and significantly differing proteins between monoculture and co-culture conditions were identified using volcano plots based on a Pearson correlation with an FDR of 0.05. Using the virulence factor database (34), Aureo Wiki (35), Uniprot (36), STRING database (37), and various studies (38–40), extracellular (ECVFs) and non-extracellular (N-ECVFs) virulence factors were identified. Results were visualized using Microsoft Excel (version 2402, build 17328.20282).

### Human oral squamous cell culture and cytotoxicity assay

The gingival squamous carcinoma Ca9-22 and buccal epithelial carcinoma HO1N1 cells were cultured in DMEM + 10% fetal bovine serum (FBS; Sigma) at 37 ⁰C, 5% CO2. Cells reached confluence within 10 days and were washed using PBS, detached by trypsinization (0.05% trypsin) for 3 minutes at 37 °C, spun down, diluted to 1•10^5^ cells/mL using DMEM/F12 (Gibco) supplemented with 10% FBS. Cells were inoculated (1•10^5^ cells/well) into wells of a 24 wells plate (Greiner) and incubated for 24 hours at 37 ⁰C, 5% CO2. Ca 9-22 and HO1N1 cells were exposed to secretome protein isolates (1:10 diluted in mDMEM-DMP, 100 mM HEPES), acquired as described above but without protease inhibitor supplementation, and incubated for 24 hours at 37 ⁰C, 5% CO2. Medium was collected and stored at -20 ⁰C until cytotoxicity assay. Cytotoxicity was determined using a LDH kit plus (Cat. No. 04 744 934 001, Roche) according to manufacturer’s protocol. Medium of the cells lysed with 1% TritonX100 in mDMEM-DMP (100 mM HEPES) was used as a positive control while the medium of unexposed macrophages was utilized as a negative control.

## Results

### Co-culturing of *C. albicans* and *S. aureus* increases the ECVF secretion by *C. albicans*

Following mass spectrometry analysis, 183 *C. albicans* proteins were detected in both samples of at least one culturing condition. To investigate the effect of *C. albicans* and *S. aureus* co-culturing on the secreted proteins, principal component analysis (PCA) was performed based on the levels of these detected proteins (**Figure 1A, B**). Based on principal component 1 (PC1, 37.5%) and PC2 (22.6%) results showed a clear separation among all groups except for *C. albicans wildtype* and *C. albicans als1/als3* ΔΔ/ΔΔ (**Figure 1A**), indicating that deletion of Als1p/Als3p does not influence *C. albicans* monoculture secretome composition. Co-culturing largely contributed to the proteins in PC1 and physical contact of *C. albicans* Als1p/Als3p to *S. aureus* contributed to the proteins in PC2. Of the 21 known *C. albicans* ECVFs (**Table S1**), 14 ECVFs were detected in both samples of at least one culturing condition and 12 ECVFs were significantly different based on ANOVA (**Figure 1C**, **Table 2**). Based on the loading plot the detected ECVFs strongly contributed to the PC1 (**Figure 1B**), suggesting that the presence of these ECVFs in the secretome strongly differs between mono and co-cultures (**Figure 1B**). Correspondingly, the presence of all detected *C. albicans* ECVFs were significantly increased in co-culture versus monoculture (**Figure 1C**, **Table 2, Figure S1**). Of these ECVFs, secreted aspartic protease 5 (Sap5) and glycosidase Crh11 were increased in all tested co-culture conditions compared to monoculture (**Figure 1C**, **Table 2**), indicating their increase was independent of Als1p/Als3p-mediated binding and other types of physical interaction. Five proteins, i.e., Plb1, Sap4, Sap6, Sod5, and Csa2, were significantly increased only in wild-type co-culture but not in als1/als3 ΔΔ/ΔΔ or transwell setting (**Figure 1C**), suggesting that their secretion was strongly dependent on Als1p/Als3p-mediated binding. The remaining five ECVFs, i.e., glycosidase Utr2, PR-1 protein homologs Rbe1 and Rbt4, Sap9, and glucan 1,3-beta-glucosidase (Xog1), were significantly increased during both regular and transwell separated co-culturing with *wildtype C. albicans*, but were not changed following co-culturing with *C. albicans* als1/als3 ΔΔ/ΔΔ (**Figure 1C**, **Table 2**), indicating their release was strongly dependent on soluble factors and independent of Als1p/Als3p-mediated binding (**Figure 1C**, **Table 2**). These significantly increased *C. albicans* ECVFs contribute to damaging functions such as proteolysis, immune evasion, cell wall modelling, and iron acquisition, suggesting that co-culturing *C. albicans* with *S. aureus* promotes the virulence potential of *C. albicans*.

**Figure 1.**
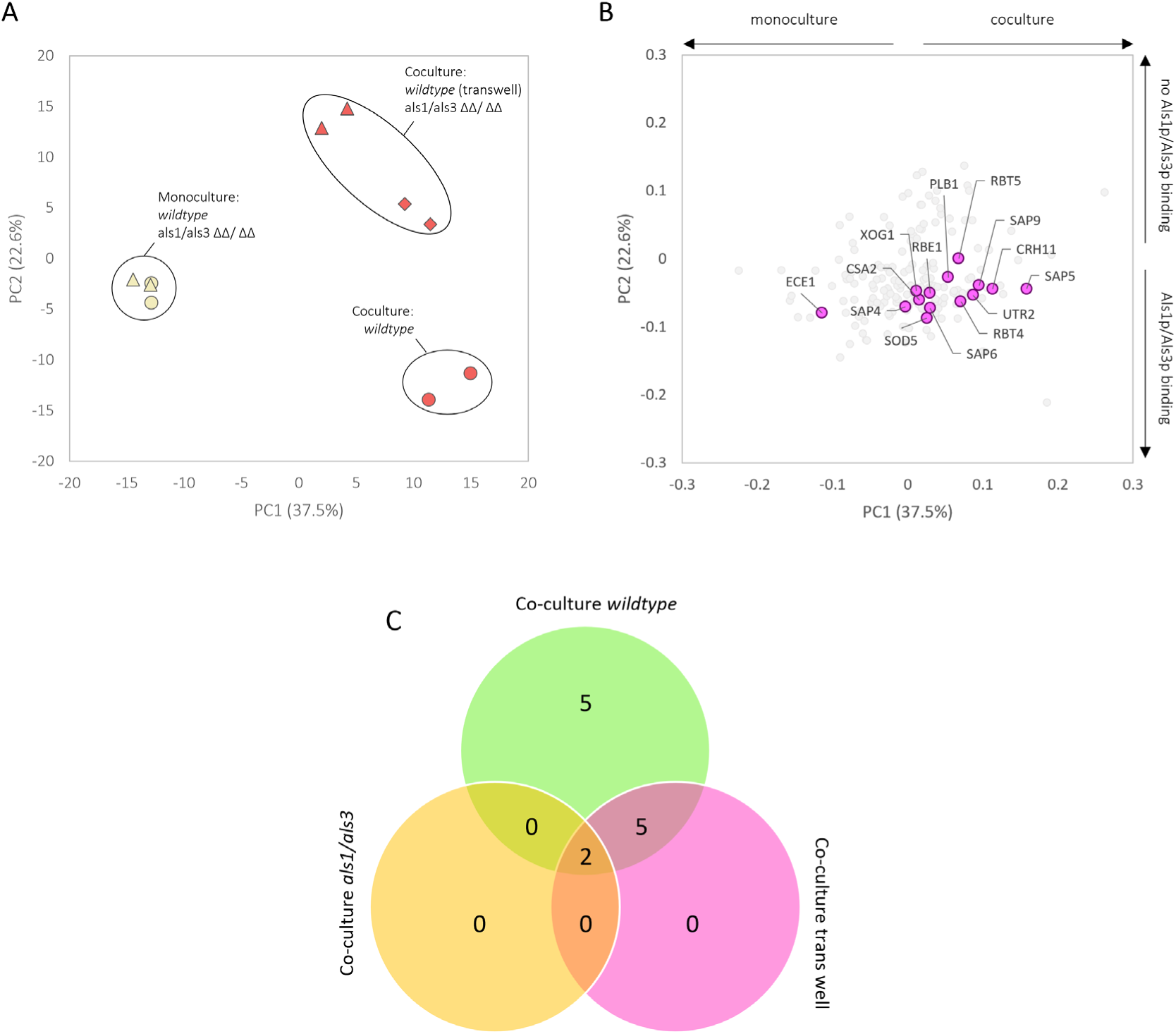
Proteomic analysis of *C. albicans* secretome revealed distinct proteins under co-culture conditions. (A) principal component analysis (yellow filled circle: *C. albicans wild type,* yellow filled triangle: *C. albicans* als1/als3 ΔΔ/ΔΔ, red filled circle: co-culture *wildtype*, red filled triangle: co-culture als1/als3 ΔΔ/ΔΔ, red filled diamond: co-culture transwell). (B) corresponding loading plot of *Candida albicans* spent medium proteins that were detectable in at both samples of at least one condition. Detected ECVFs are coloured in pink. (C) Venn diagram of significantly differing (ANOVA) ECVF proteins in the secretome of *wildtype*, trans well, and als1/als3 ΔΔ/ΔΔ co-cultures.

**Table 2.** ***C. albicans* ECVFs are mainly increased due to Als1p/Als3p binding by *S. aureus*.** Significantly differing *C. albicans* ECVF proteins in the secretome of *wildtype*, trans well, and als1/als3 ΔΔ/ΔΔ co-cultures. All separated by regulation, together with corresponding virulence class, protein description, gene, and log 2-fold change per co-culture condition.

| Regulation | Virulence class | Description | Gene | Log 2-fold change |  |  |
| --- | --- | --- | --- | --- | --- | --- |
| Regulated by the presence of *S. aureus* ●●● |  |  |  | Co-culture wildtype | Co-culture als1/als3 | Co-culture trans well |
| up | Cell wall remodelling | Glycosidase | Crh11 | 2,94 | 0,35 | 1,68 |
| up | Enzyme (proteolytic) | Secreted aspartic protease 5 | Sap5 | 2,72 | 1,01 | 1,46 |
| Regulated by Als1p/Als3p binding to soluble *S. aureus* factors ●● |  |  |  |  |  |  |
| up | Cell wall remodelling | Glycosidase | Utr2 | 2,93 | ns | 1,40 |
| up | Enzyme (proteolytic) | Secreted aspartic protease 9 | Sap9 | 1,85 | ns | 0,94 |
| up | Immune evasion | PR-1 proteins homologue | Rbt4 | 2,91 | -0,23 | 1,11 |
| up | Immune evasion | PR-1 proteins homologue | Rbe1 | 1,71 | -1,63 | 1,67 |
| up | Immune evasion | Glucan 1,3-beta-glucosidase | Xog1 | 1,40 | -2,00 | 0,82 |
| Regulated by physical binding of Als1p/Als3p to *S. aureus* ● |  |  |  |  |  |  |
| up | Enzyme (lipolytic) | phospholipase 1 | Plb1 | 2,16 | ns | ns |
| up | Enzyme (proteolytic) | Secreted aspartic protease 4 | Sap4 | 1,85 | -0,90 | -0,66 |
| up | Enzyme (proteolytic) | Secreted aspartic protease 6 | Sap6 | 2,45 | -0,51 | -0,17 |
| up | Immune evasion | Superoxide dismutase 5 | Sod5 | 1,73 | -1,60 | ns |
| up | Iron utilization | Secreted hemophore | Csa2 | 2,18 | -1,42 | ns |

### Co-culturing of *C. albicans* and *S. aureus* promotes ECVF secretion by *S. aureus*

Following mass spectrometry analysis, 930 *S. aureus* proteins were detected in at least one culturing condition. Principle component analysis indicates a clear separation between monocultures and co-cultures, as well as between unbuffered and buffered *S. aureus* monocultures (**Figure 2A**), suggesting that both medium buffering and the presence of *C. albicans* have substantial yet distinct effects on the proteins secreted by *S. aureus*. It is worth noting that no separation was observed among three co-culture conditions (**Figure 1A**), indicating that deletion of Als1p/Als3p or lack of physical interaction with *C. albicans* do not influence the *S. aureus* secretome when compared to wildtype co-culture. Of the 50 known *S. aureus* ECVFs (**Table S2**), 27 were detected in both samples of at least one culture condition and 20 were found statistically significantly changed (**Figure 2B-C**, **Table 3**). Detected *S. aureus* ECVFs contributed more strongly to PC1 (43%) than PC2 (28.7%) (**Figure 2B**), indicating that their presence is mainly affected by co-culturing with *C. albicans*. Due to the clustering of both *wildtype* and als1/als3 ΔΔ/ΔΔ co-culture conditions during PCA, the effect of co-culturing is likely Als1p/Als3p independent. Of the detected ECVFs alpha haemolysin (Hly/Hla), beta haemolysin (Hlb), and gamma haemolysin gamma (HlgA-C) were increased in all conditions (**Table 3, Figure S2**).

**Figure 2.**
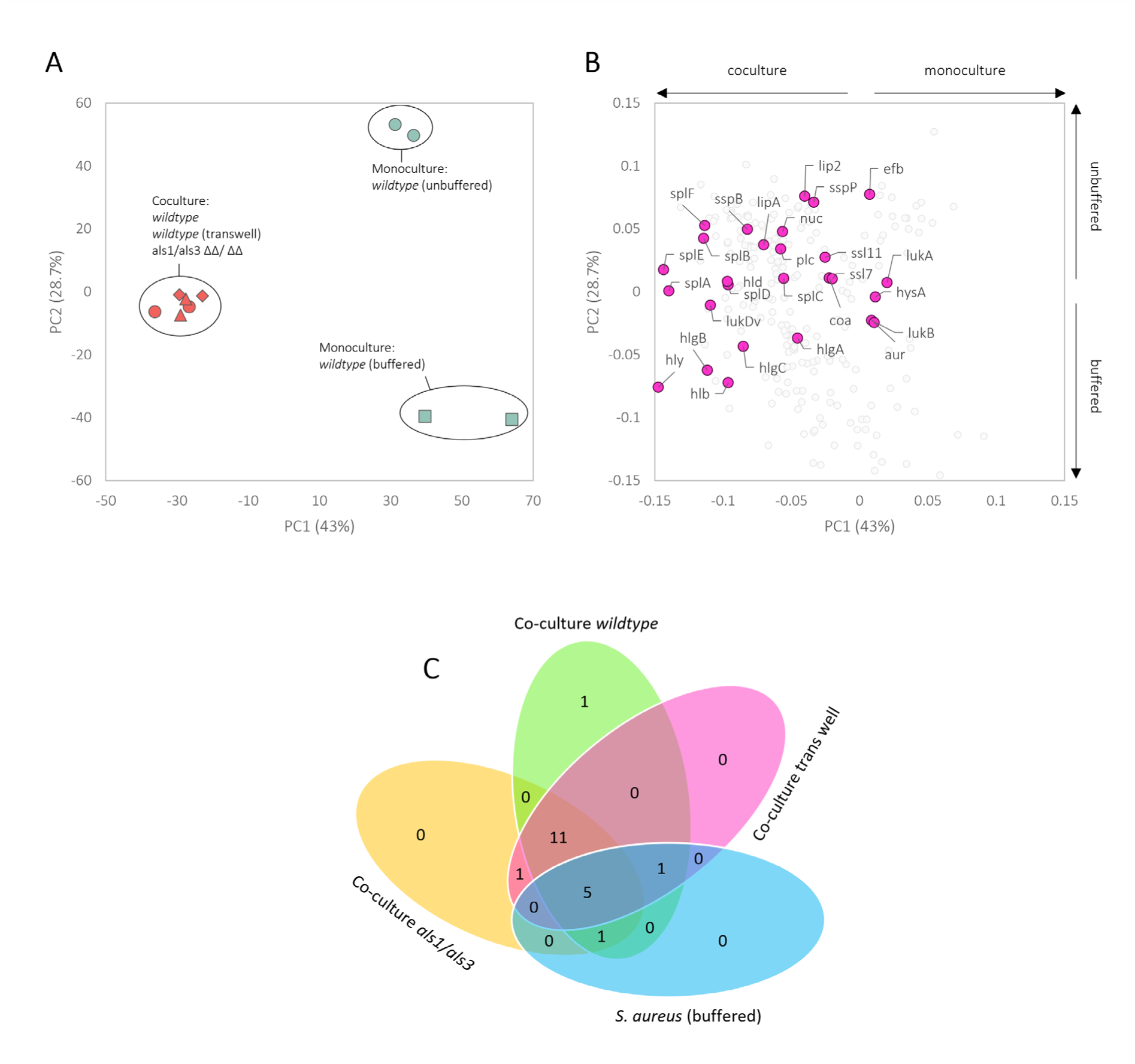
Proteomic analysis of *S. aureus* secretome revealed distinct proteins under co-culture and buffered conditions. (A) principal component analysis and (B) corresponding loading plot of *Staphylococcus aureus* spent medium proteins that were detectable in both samples of at least one condition. ECVFs are coloured in pink. (C) Venn diagram of significantly differing ECVF proteins in the secretome of *wildtype*, trans well, and als1/als3 ΔΔ/ΔΔ co-cultures as well as buffered monoculture.

**Table 3.**
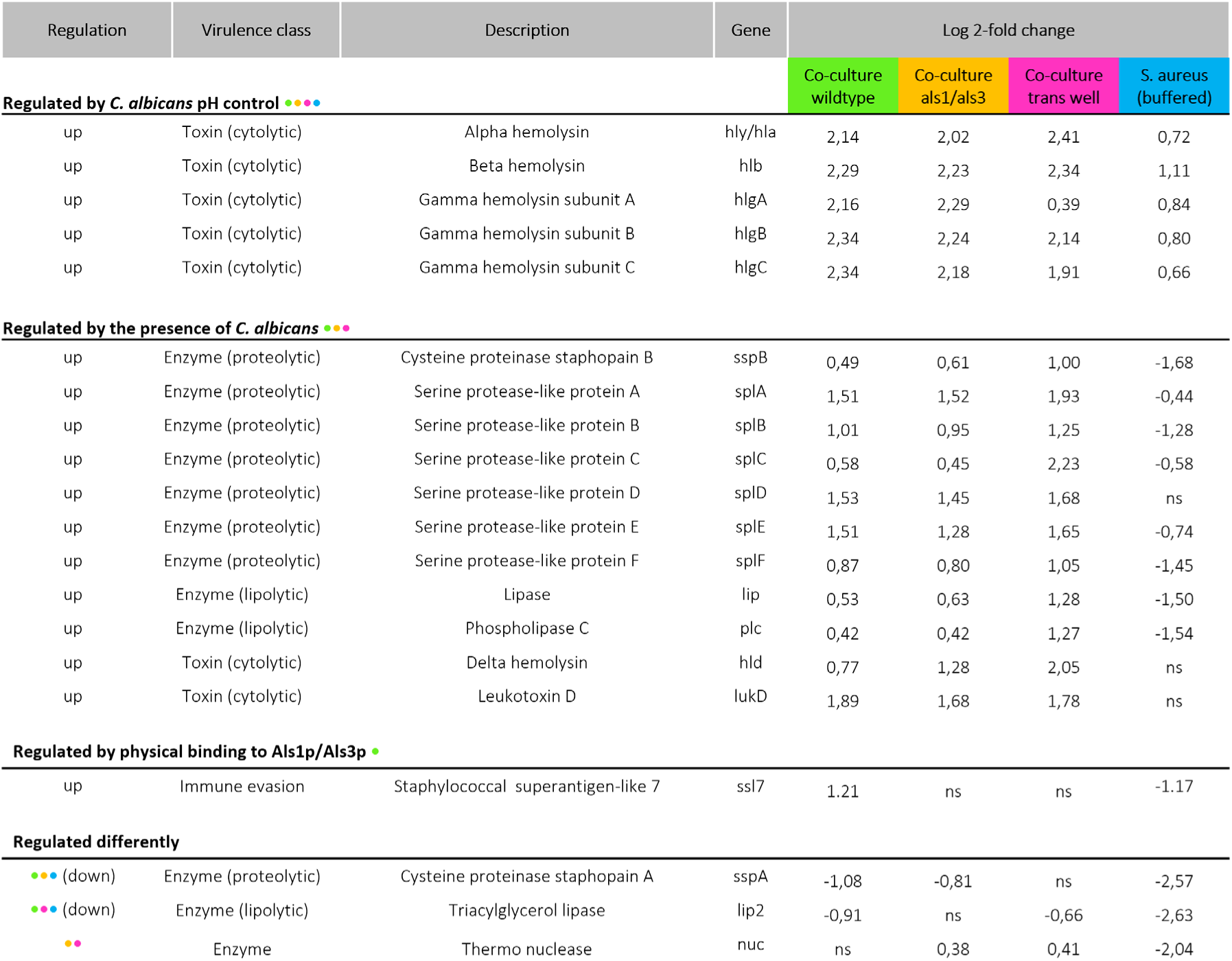
***S. aureus* ECVFs are mainly increased by the presence of *C. albicans* and its pH maintenance.** Significantly differing *S. aureus* ECVF proteins in the secretome of *wildtype*, trans well, and als1/als3 ΔΔ/ΔΔ co-cultures as well as buffered monocultures. All separated by regulation, together with corresponding virulence class, protein description, gene, and log 2-fold change per culture condition. ns: not significant?

Buffering the pH alone increased the secretion of these proteins, and this increase was maintained despite the lack of physical contact or Als1p/Als3p-mediated binding. This results, together with the high pH during *C. albicans-S. aureus* c o-culture (**Figure S3**), suggests that *C. albicans*-maintained pH likely contributed to the increase of the five cytolytic haemolysin proteins. Cysteine proteinase staphopain B (SspB), serine protease-like protein A-F (SplA-F), lipase (Lip), phospholipase C (Plc), delta haemolysin (Hld), and leukotoxin D (LukD) were significantly more present in all co-culture conditions, but significantly less present or unaltered during buffered monoculturing (**Table 3**).

Because Als1p/Als3p deletion and separated growth did not deviate from *wildtype* co-culture results, these proteins were increased due to the presence of *C. albicans*. Contrasting to *C. albicans* ECVFs, *S. aureus* ECVF composition was mildly affected by Als1p/Als3p binding. Only staphylococcal superantigen-like 7 (Ssl7) was significantly higher in *wildtype* co-culture while not significantly affected during the other tested co-culture conditions and decreased during buffered monoculture growth (**Table 3**).

Altogether, co-culturing of *C. albicans* and *S. aureus* promotes the secretion of ECVFs. Whereas the secretion of *C. albicans* virulence factors is mainly regulated by Als1p/Als3p binding, the majority of *S. aureus* ECVFs are regulated by the presence of *C. albicans* or its pH maintenance. The greater part of ECVFs that are upregulated during co-culturing (*C. albicans*: 5 out of 11, *S. aureus*: 16 out of 21) were classified as either cytolytic, proteolytic, or lipolytic (**Table 2, 3)**, indicating that the damaging properties of the secretome increases during co-culturing.

### Co-culturing increases the levels of non-extracellular virulence factors from both *C. albicans* and *S. aureus*

To identify other potential virulence factors that might contribute to the elevated cytotoxicity, we identified 42 non-extracellular virulence factors (N-ECVFs) that are not known to be actively secreted. Of these 42 N-ECVFs, 14 belong to *C. albicans* and mainly contributed to co-culture (PC1, 37.5%) and Als1p/Als3p deletion (PC2, 22.6%) based separation (**Figure 3A**). All of the 14 identified *C. albicans* N-ECVFs were significantly changed in co-cultures compared to monocultures (**Figure 3B**). Level of hyphae-regulated cell wall protein 1 (Hyr1) and heat shock protein 70 (Ssa1) were significantly increased in *wildtype* co-culture versus *C. albicans* monoculture, but this change was attenuated in als1/als3 ΔΔ/ΔΔ co-culture and transwell setting (**Figure 3C**). Additionally, the level of thioredoxin peroxidase (Ahp1) and Glycolipid 2-alpha-mannosyltransferase 1 (Mnt1) were significantly decreased in *wildtype* and transwell co-culture, and this decrease was enhanced in als1/als3 ΔΔ/ΔΔ co-culture. These results suggest that Hyr1, Ssa1, Ahp1, and Mnt1 are influenced by both Als1p/Als3p-dependent *C. albicans-S. aureus* interaction and other *S. aureus* factors. The level of cell wall protein (Rhd3) and cell wall remodelling related proteins Ihd1, Sun41, Pga4, Als1p, Als3p, Mp65, and Bgl2 were significantly increased in wildtype co-culture (**Figure 3C**). This increase was attenuated by the physical separation, reflected by less increase in transwell co-culture, and completely abolished in als1/als3 ΔΔ/ΔΔ co-culture (**Figure 3C**), indicating the cell wall remodelling was strongly influenced by Als1p/Als3p-mediated binding to *S. aureus*. Glutathione reductase (Glr1) presence was only increased during co-culturing with *wildtype C. albicans* (**Figure 3C**) and, therefore, regulated by the physical binding of *S. aureus* to Als1p/Als3p. Finally, the enolase (Eno1) level was not altered during wildtype co-culturing, while its level was decreased in als1/als3 ΔΔ/ΔΔ and trans well co-cultures (**Figure 3C**). Most of the 14 significantly changed *C. albicans* N-ECVFs were related to adhesion or cell wall remodelling, suggesting that physical interaction between *C. albicans* and *S. aureus* promotes the levels of these virulence factors.

**Figure 3.**
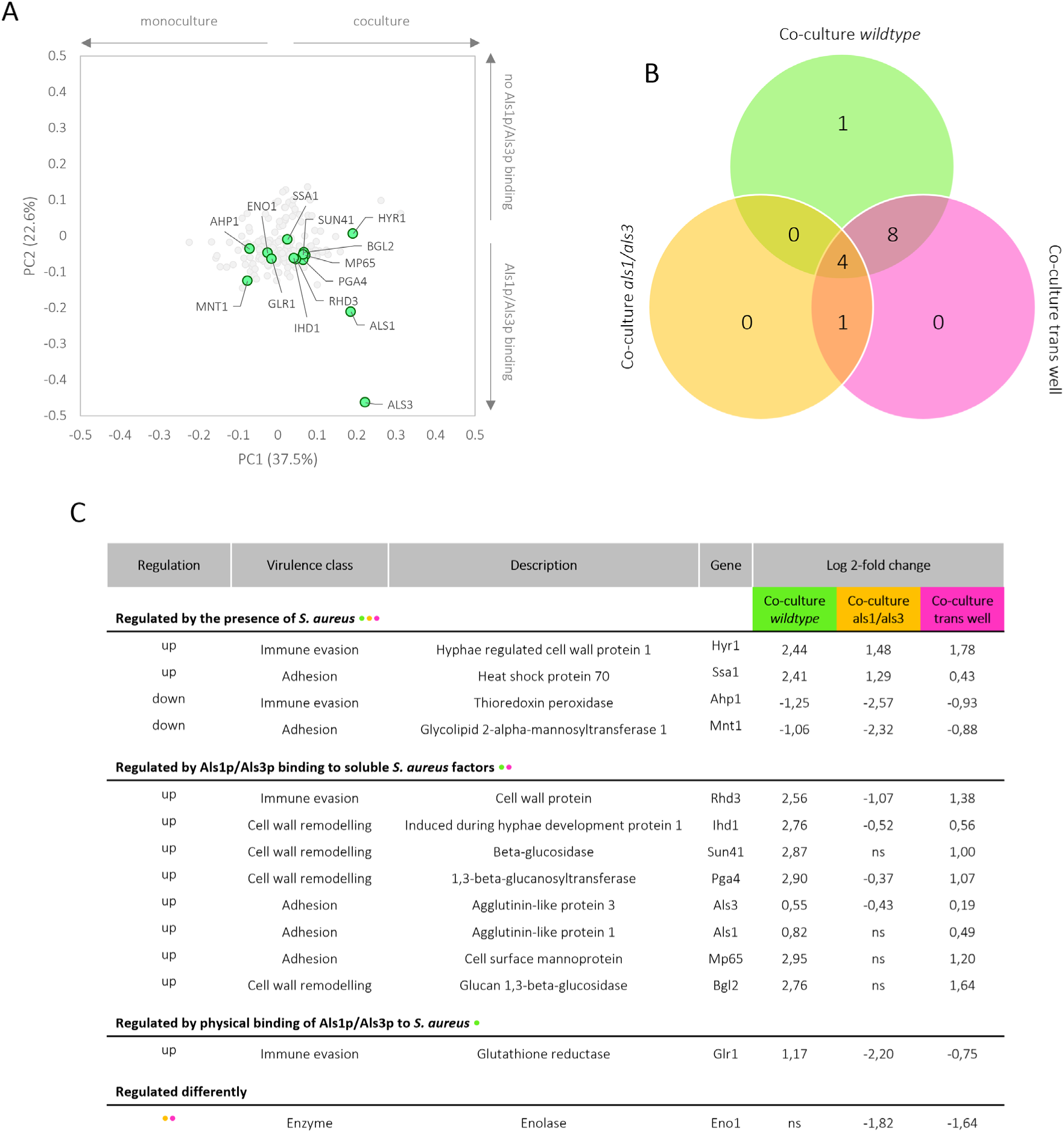
Proteomic analysis of *C. albicans* secretome revealed distinct N-ECVF proteins under co-culture conditions. (A) score plot, corresponding to the PCA analysis performed in Figure 1A, depicting *C. albicans* N-ECVFs that were identified in both samples of at least one culture condition (green). (B) Venn diagram of significantly differing N-ECVF proteins in the secretome of *wildtype*, trans well, and als1/als3 ΔΔ/ΔΔ co-cultures. (C) The table containing all significantly differing *C. albicans* N-ECVF proteins in the secretome of all tested conditions. All are separated by regulation, together with corresponding virulence class, protein description, gene, and log 2-fold change per culture condition.

### *S. aureus* non-extracellular virulence factors

The 28 identified *S. aureus* N-ECVFs mainly contributed to co-culture secretome separation (PC1 (43%)) but both buffered and unbuffered conditions (PC2 (28.7%)) (**Figure 4A**). Of the 28 identified *S. aureus* N-ECVFs, 17 were significantly different based on ANOVA (**Figure 4B**). Buffering strongly impacted N-ECVFs levels, evidenced by the fact that 11 and 2 N-ECVFs in buffered monoculture were significantly higher and lower than those in unbuffered monoculture, respectively (**Figure 4C**). This pH-mediated effect seemed to be divergently interfered with by other factors from *C. albicans*. For example, the buffering-increase of d-alanine-d-alanyl carrier protein ligase (DltA) and enolase (Eno) was further enhanced by the presence of *C. albicans,* and this enhancement is independent of Als1p/Als3p or physical proximity (**Figure 4C**). For those proteins that were decreased by buffering, i.e., ClfB, Ebp, Oata, Srap/Sasa, IsdA, IsdC, IsdE, SpA, SdrC, and ClfA, the decrease was attenuated or even reserved by the presence of *C. albicans*, despite that *C. albicans* maintained pH comparably to HEPES buffering. These results suggest that other *C. albicans*-derived soluble factors have stronger effects in influencing these proteins compared to pH. Interestingly, the deletion of Als1p/Als3p had mild effects on the N-ECVFs of *S. aureus*. Only extracellular adherence protein/MHC analogous protein (Eap/Map) and iron-regulated surface determinant (IsdB) exhibited different trends in both co-cultures (**Figure 4C**). Together, these results suggest that both *C. albicans*- derived pH maintenance has a broader impact than that of Als1p/Als3p-mediated binding on N- ECVFs of *S. aureus*.

**Figure 4.**
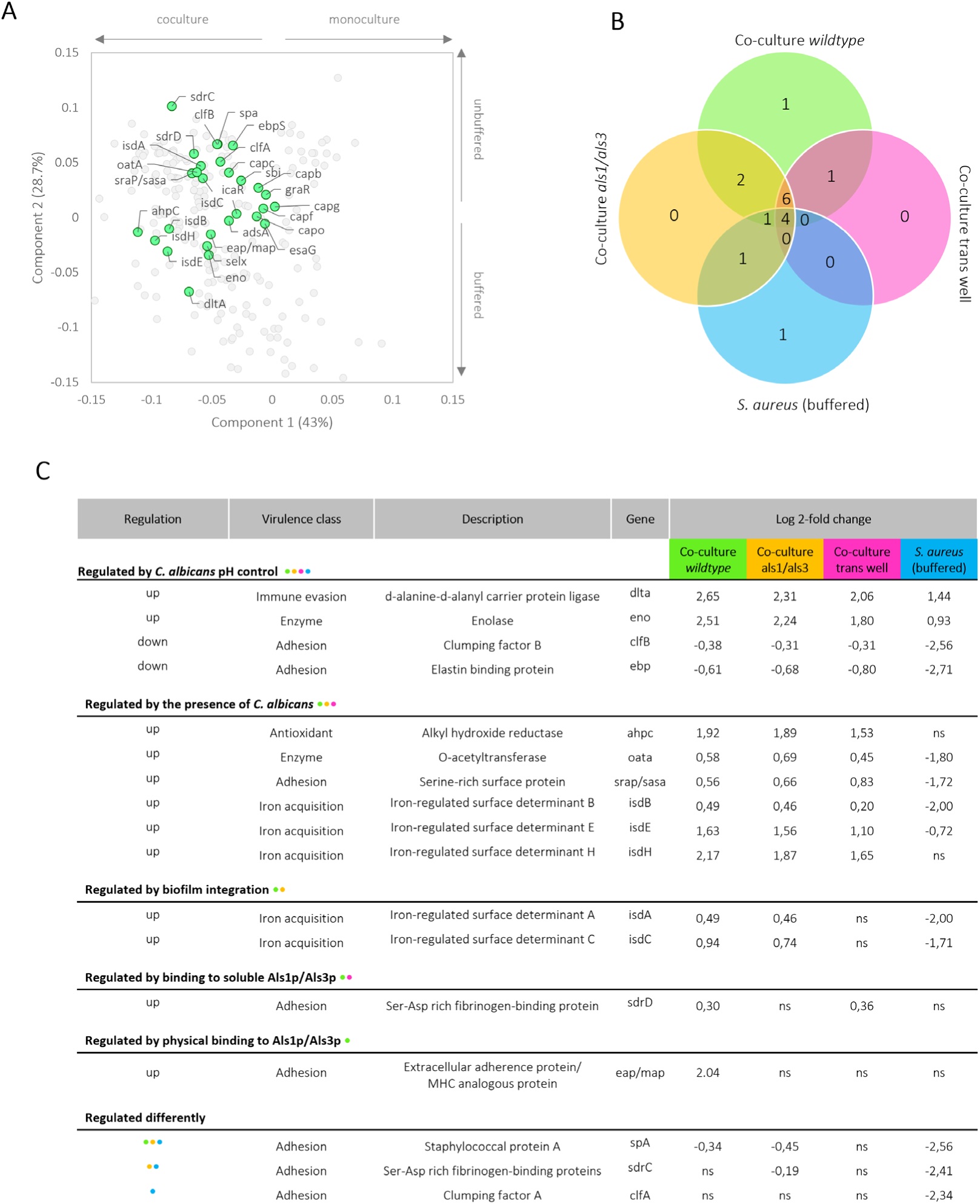
**Proteomic analysis of *S. aureus* secretome revealed distinct N-ECVF proteins under co-culture conditions. (**A) score plot, corresponding to the PCA analysis performed in Figure 1A, depicting *S. aureus* N-ECVFs that were identified in both samples of at least one culture condition (green). (B) Venn diagram of significantly changed N-ECVFs in the secretome of *wildtype*, trans well, and als1/als3 ΔΔ/ΔΔ co-cultures as well as buffered monoculture. (C) The table containing all significantly differing *S. aureus* N-ECVFs in the secretome of all tested conditions. All are separated by regulation, together with corresponding virulence class, protein description, gene, and log 2-fold change per culture condition.

### Co-culturing promotes cytotoxicity of both *C. albicans* and *S. aureus* to oral squamous cells

Reasoning that most damaging ECVFs of *C. albicans* and of *S. aureus* wildtype co-cultures was relatively higher than monoculture (**Figure S1A and S1B**), and many N-ECVFs were increased during coculturing (**Figure 4C**), we hypothesized that the secretomes of *C. albicans*-*S. aureus* co-cultures induces greater damage to host cells compared to monoculture. To test this, we exposed human gingival squamous Ca 9-22 and human buccal mucosa squamous HO1N1 cells to undiluted mono and co-culture secretomes of *C. albicans wildtype, C. albicans* als1/als3 ΔΔ/ΔΔ, and/or *S. aureus* (cultured in either buffered or unbuffered medium). To evaluate cytotoxicity, medium lactate dehydrogenase activity was measured following the 24 hour exposure period. All monoculture secretomes, i.e., *C. albicans wildtype, C. albicans* als1/als3 ΔΔ/ΔΔ, and *S. aureus* (unbuffered), induce similar levels of cytotoxicity as the negative control (∼20-30%, **Figure S4**). In contrast, all tested co-culture secretomes and *S. aureus* buffered induced higher cytotoxicity to Ca 9-22 cells (**Figure S4C**). Similarly, the co-culture secretomes induced higher cytotoxicity to HO1N1 cells (**Figure S4D**). These results are in agreement with the observed elevated levels of ECVFs in co-culture secretomes compared to monoculture. Interestingly, the secretome of buffered *S. aureus* monocultures showed higher cytotoxicity to Ca 9-22 (92%) and HO1N1 (79%) (**Figure S4C and S4D**), despite that the levels of ECVFs in buffered *S. aureus* are comparable or lower than that in co-cultures (**Table 3**). These results suggest that the cytolytic *S. aureus* ECVFs, promoted by *C. albicans* pH maintenance, are likely accountable for the increased cytotoxicity during the co-culture.

## Discussion

Previous studies have shown that co-infections of *C. albicans* and *S. aureus* significantly promote lethality compared to mono infections (14–22,41). Co-invasion and dissemination of *S. aureus* are crucial to this process and are facilitated by the secretion of damaging ECVFs and hyphal invasion of *C. albicans*. In this study, we applied proteomics to elucidate the ECVFs and N-ECVFs released by both *C. albicans* and *S. aureus*. Our results showed that co-culturing significantly increased the levels of ECVFs and N-ECVFs. Whereas Als1p/Als3p binding mainly influenced the virulence factors released by *C. albicans*, *C. albicans*-derived pH maintenance primarily contributed to the increase of virulence factors released by *S. aureus*. The increase in virulence factors coincided with the increase in cell cytotoxicity toward human oral cells.

### *C. albicans* virulence is promoted by *S. aureus* during co-culturing and mainly attributed to Als binding

ECVFs are essential for various pathogenic processes of *C. albicans,* such as hyphal formation, damaging host cells, invasion, and immune evasion. Previously, Als3, Csa2, Rbt4, Sap4 and Sap6 proteins were found to be enriched in the secretome of N-acetylglucosamine-induced hyphal growth over yeast (40,42–46) and contribute to the virulence of *C. albicans* pathogenesis (40,42–46). Deletion of *rbt4* (40,42–46)*, sod5* (46), *sap4*, and *sap6* (47), significantly attenuates or completely diminishes *C. albicans* lethality in animal models. Mechanistically, double deletion of *rbt4* increased sensitivity to attack by polymorphonuclear leucocytes (40,42–46), and deletion of *sap4-6* increased the sensitivity to macrophage-mediated killing (48). Similarly, we showed that hyphal growth of *C. albicans* was promoted in co-culture with *S. aureus* (data not shown), and Rbt4, Sap4-6, Sod5, and Csa2 were increased in co-culture. Consistently, transcription of *Sod5* was increased in hyphal growth (40,42– 46), and Sod proteins are known to aid in the protection against intracellular neutrophil killing (46). In addition to these proteins, we found that hyphae-related proteins (49,50), such as Als1, Als3, Ihd1, and Hyr1, are higher in co-cultures over monoculture. Furthermore, the co-culture promotes the level of Xog1 in the secretome. Xog1 is an exo-1,3-beta-glucanase essential for reducing beta-glucan epitope exposure (β-glucan masking) to immune cells, hence reducing the phagocytotic interaction and enhancing immune evasion (51) (52). Other two cell wall crosslinking enzymes Crh11 and Utr2 were also upregulated during co-culturing. Crh11 and Utr2 were shown to impact β-glucan masking mildly (52). Together, these results support the contribution of these proteins in *C. albicans-S. aureus* co-culture to the pathogenesis during co-infection. Apart from *S. aureus*, *S. epidermidis* and mitis group streptococci such as *S. gordonii*, *S. oralis*, and *S. sanguinis* bind *C. albicans* hyphae through interactions with Als3p (53–56), suggesting that these bacteria may provoke similar effects. Nevertheless, of all these bacteria, *S. aureus* has been shown to bind hyphae best (57), indicating that *S. aureus* will likely alter proteins regulated by Als1/Als3 binding more than the other bacteria.

### *S. aureus* virulence is significantly promoted by *C. albicans* during co-culturing

Similarly to *C. albicans*, the co-culturing also promotes the virulence potential of *S. aureus* by increasing the secretary level of cytolytic, proteolytic, or lipolytic proteins. We found *C. albicans-* mediated pH maintenance is a critical factor in regulating these proteins. This is consistent with previous observations, where *C. albicans* tended to maintain a neutral pH during co-culturing (**Figure S3**) (25) and, thereby, promoted the production and secretion of alpha hemolysin (hla) (25). In addition to hla, we also found beta and gamma hemolysins to be increased in the presence of *C. albicans*. Mechanistic studies showed that the P3 promoter of the *S. aureus agr* system was activated by *C. albicans,* which increases the expression of RNAIII, essential for the production and secretion of these toxins (58,59). While we and others confirmed the contribution of pH in regulating *S. aureus* virulence factors, other *C. albicans*-derived factors contribute to the elevated virulence potential. For example, the ECVFs were significantly decreased in buffered monocultures over unbuffered monoculture but significantly increased in co-culture over unbuffered monoculture. Therefore, these virulence factors are likely affected by both pH maintenance and other *C. albicans* factors. These results further highlighted the nuanced regulation of the secreted virulence factors by chemical and biological cues during the co-culture.

### *C. albicans* and *S. aureus* reciprocally promote the iron acquisition potential of each other

An important strategy for pathogenic microbes is to exploit host haem/iron sources during the infection. For *C. albicans*, Csa2 is involved in the uptake of haemoglobin and haem proteins and their utilization as an iron source (43) and Csa2-mediated haemoglobin-iron acquisition is assisted by Rbt5, Pga7, Frp1, and Frp2 (60–62). We found that co-culturing increases the secreted level of Csa2, but not Rbt5, Pga7, Frp1, and Frp2 in *C. albicans*. In contrast to *C. albicans*, we found co-culturing has a broader impact on the haemolytic/iron acquisition potential of *S. aureus*. It is well established that Isd (iron-regulated surface determinant) system are co-ordinately involved in haemoglobin and heme binding, uptake, and iron release (63). Of all the isd proteins, we found that the levels of cell wall anchored isdA, isdB, isdC, isdE, and isdH, but not intracellular isdI and isdG, were significantly higher in co-culture. This is partially inconsistent with a proteomic analysis on whole-cell proteins in co-culture over monoculture, where it was found that none of these protein significantly changed in co-culture (57). These results suggest that the release of cell wall anchored isd proteins was increased during co-culture. Further analysis warranted the mechanisms underlying this effect.

### Co-culturing increases the cytotoxicity

As many virulence factors from both *C. albicans* and *S. aureus* were dramatically increased in co-culturing, it is not surprising that the secretome from co-culture exhibited higher cytotoxicity toward Ca 9-22 and HO1N1 cells compared to that from the monocultures. While our results need to be repeated Consistently, a similar increase of cytotoxicity was observed for *C. albicans-S. aureus* co-culturing toward keratinocytes NOK-si and HaCat cells, although the extent of cytotoxicity increase differed (24). Nevertheless, we found secretome from *C. albicans* monoculture has negligible cytotoxicity to both Ca 9-22 and HO1N1 lines, which is inconsistent with the high toxicity observed using keratinocytes NOK-si and HaCat cells (24). This discrepancy might be due to the different sensitivity of these cells. In addition to *C. albicans*, *S. aureus* contributed to the cytotoxicity induced by secretome from co-cultures. Surprisingly, the cytotoxic effect of the secretome from buffered *S. aureus* monoculture was higher than all co-culture secretomes. While the exact reasons are hard to decipher, this might be because of a higher amount of total protein in buffered *S. aureus* monoculture secretomes than in all other tested conditions (**Figure S3E**). Interestingly, co-culture biofilms contained similar amounts of *S. aureus* DNA to buffered *S. aureus* monoculture biofilms (**Figure S3B**). This result and the notion that most elevated virulence factors are secretory proteins suggest that the presence of *C. albicans* promoted an active secretion by *S. aureus*.

## Acknowledgements

This work was supported by the University of Amsterdam Research Priority Area Systems Biology Host–Microbiome Interactions. S.B. is supported by the University of Amsterdam Centre for Urban Mental Health. Finally, S.B., B.P.K. and J.Z. are supported by NWA-ORC project 1389.20.080. We thank Caroline Bosch-Tijhof and Caroline de Jongh (ACTA) for the cultivation and testing of the Ca 9-22 and HO1N1 cell lines, Winfried Roseboom (UvA) for assistance on LC-MS/MS analysis. We thank dr. ing. Girbe Buist (University of Groningen) for kindly providing the *S. aureus* agr strains. We thank Dennis Rijnsburger, Pim van Leeuwen, Marco Lezzerini, and dr. Belinda Koenders-van Sintanneland for general support and lab management.

## Contributions (CRediT author statement)

Conceptualization: R.P., J.Z.; Methodology, Investigation: R.P.; Formal analysis, Data Curation: R.P., G.K.; Visualization: R.P.; Writing - Original Draft: R.P.; Supervision: B.P.K., S.B., S.A.J.Z. ; Writing - Review & Editing: B.P.K., S.B., S.A.J.Z., J.Z.; Funding acquisition: B.P.K., S.B., J.Z.

