## Supplementary file for "*Candida albicans* and *Staphylococcus aureus* reciprocally promote secretion of virulence factors"

**Table S1.** Known *C. albicans* ECVFs with corresponding gene and possible damaging potential.

| Virulence factor | Gene | Damaging potential |
|---|---|---|
| ***C. albicans* damaging ECVFs** | | |
| Candidalysin | Ece1 | Cytolytic |
| Secreted aspartic protease 1-10 | Sap1-10 | Proteolytic |
| Lysophospholipase 1 | Plb1 | Lipolytic |
| ***C. albicans* non-damaging ECVFs** | | |
| Glycosidase CRH11/CRH12 | CRH11/12 | |
| PR-1 proteins homologue | RBE1 | |
| Glycosidase UTR2 | UTR2 | |
| PR-1 proteins homologue | RBT4/5 | |
| Superoxide dismutase 5 | SOD5 | |
| Secreted hemophore CSA2 | CSA2 | |
| Glucan 1,3-beta-glucosidase | Xog1 | |

**Table S2.** Known *S. aureus* ECVFs with corresponding gene and possible damaging potential.

| Virulence factor | Gene | Damaging potential |
|---|---|---|
| ***S. aureus* damaging ECVFs** | | |
| Aureolysin | aur | Cytolytic |
| Alpha hemolysin | hly/hla | Cytolytic |
| Beta hemolysin | hlb | Cytolytic |
| Delta hemolysin | hld | Cytolytic |
| Gamma hemolysin | hlgA, B, C | Cytolytic |
| Leukotoxin A, B, D, E | lukA, B, D, E | Cytolytic |
| Serine protease | sspA | Proteolytic |
| Cysteine protease | sspB | Proteolytic |
| Serine protease | splA-F | Proteolytic |
| Lipase | geh | Lipolytic |
| Triacylglycerol lipase | lip2 | Lipolytic |
| Phospholipase C | plc | Lipolytic |
| ***S. aureus* non-damaging ECVFs** | | |
| Clumping factor A, B | clfA, B | |
| Collagen adhesion | can | |
| Elastin binding protein | ebp | |
| Extracellular adherence protein/MHC analogous protein | eap/map | |
| Fibrinogen binding protein | efb | |
| Fibronectin binding proteins | fnbA, B | |
| Intercellular adhesin | icaA-D, R | |
| Ser-Asp rich fibrinogen-binding proteins | sdrC, D, H | |
| Staphylococcal protein A | spa | |
| Staphylococcal superantigen-like 1-11 | ssl1-11 | |

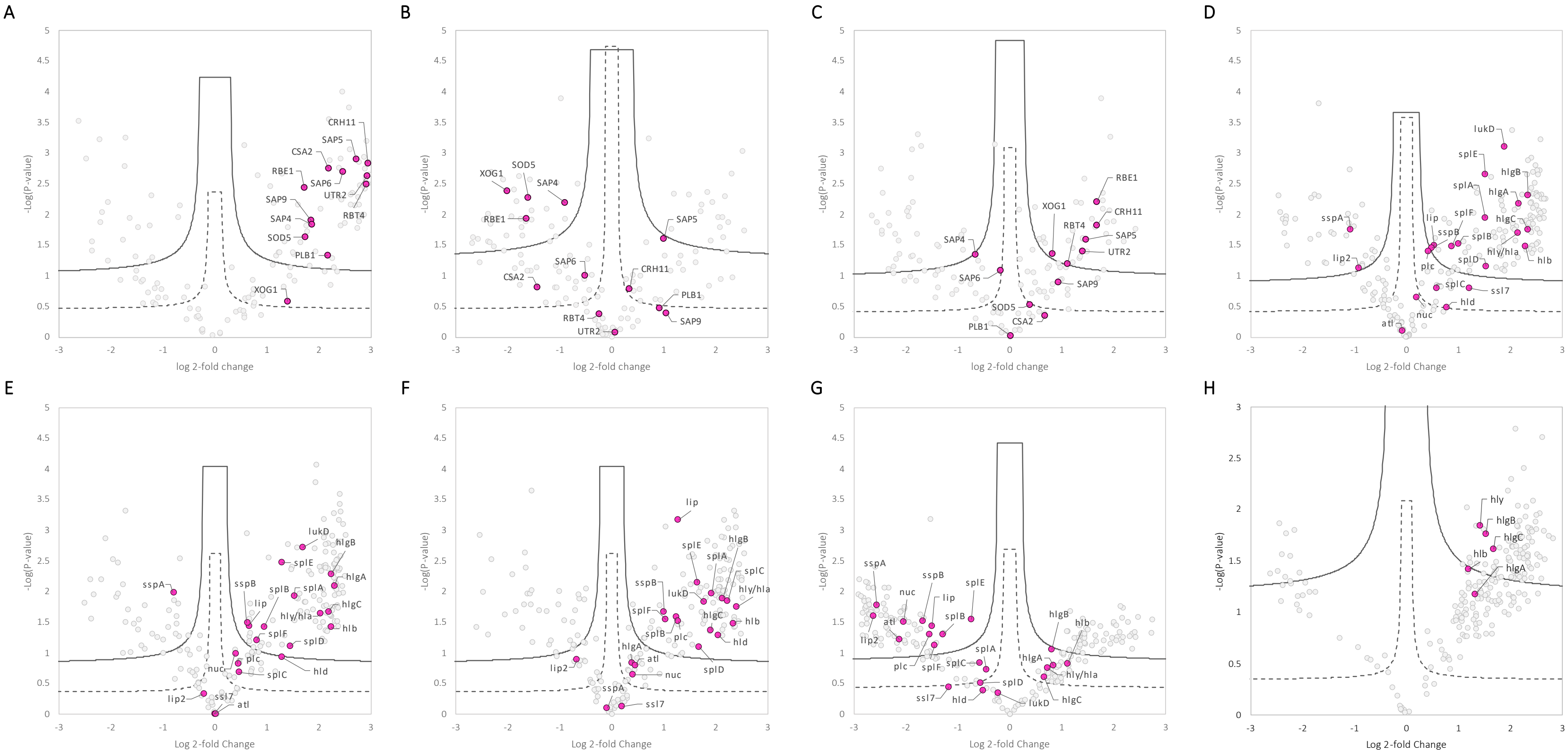


**Figure S1 Difference in secretomal protein presence, based on Log2 fold changes and -log p-value** (A) co-culture *wildtype* versus *C. albicans wildtype* monoculture, (B) co-culture als1/als3 ΔΔ/ΔΔ versus *C. albicans* als1/als3 ΔΔ/ΔΔ monoculture, (C) co-culture *wildtype* (trans well) versus *C. albicans wildtype* monoculture. (D) co-culture *wildtype* versus *S aureus* monoculture (unbuffered), (E) co-culture als1/als3 ΔΔ/ΔΔ versus *S aureus* monoculture (unbuffered), (F) co-culture *wildtype* (trans well) versus *S aureus* monoculture (unbuffered), (G) *S aureus* monoculture (100 mM HEPES) versus *S aureus* monoculture (unbuffered). (H) co-culture *wildtype* versus *S aureus* monoculture (100 mM HEPES). ECVFs are coloured pink and non ECVF proteins light grey. Significance lines are based on either a FDR of 0.05 (dashed) or 0.01 (full line). Proteins above the FDR 0.05 line were considered significantly differing.

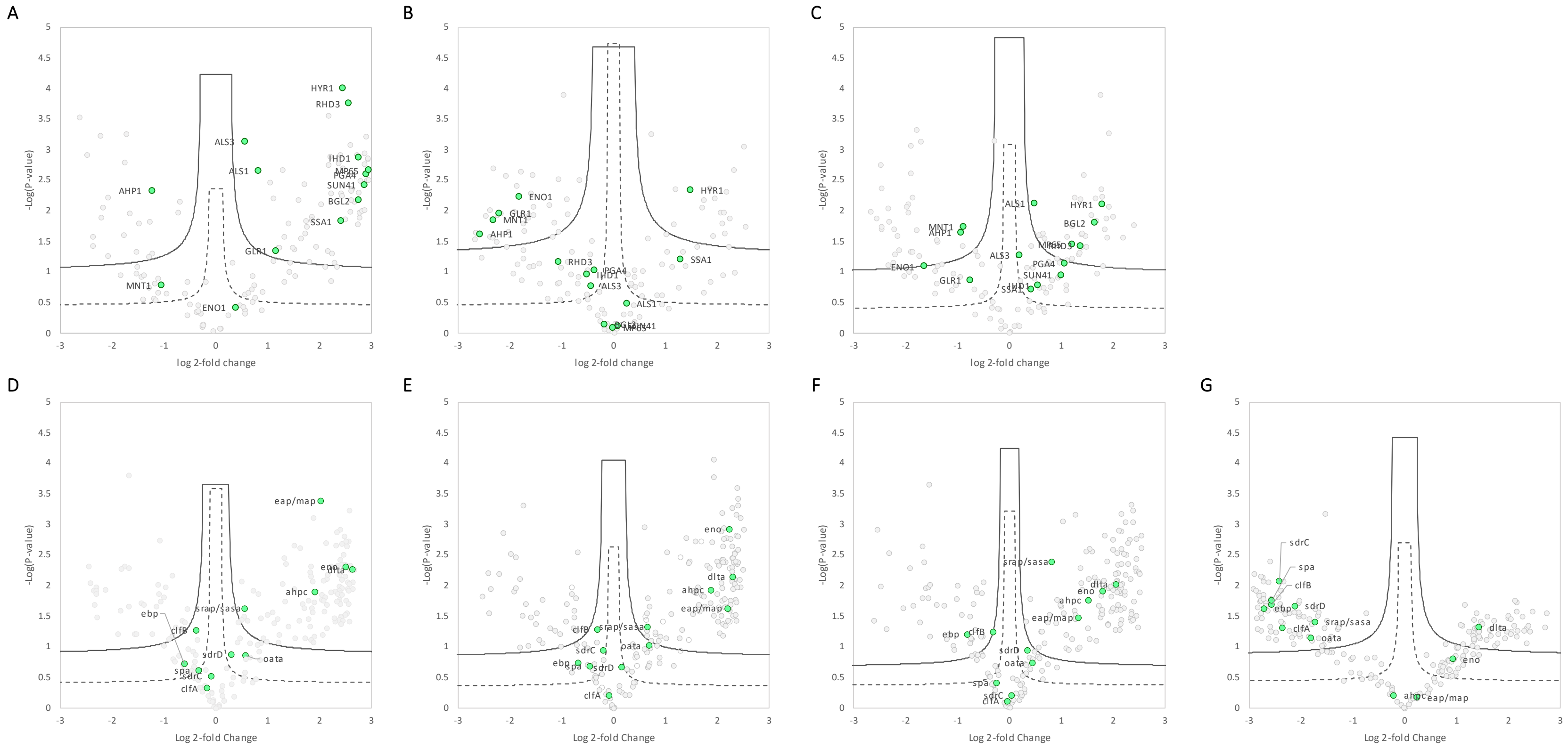

**Figure S2 Difference in secretomal protein presence, based on Log2 fold changes and -log p-value** (A) co-culture *wildtype* versus *C. albicans wildtype* monoculture, (B) co-culture als1/als3 ΔΔ/ΔΔ versus *C. albicans* als1/als3 ΔΔ/ΔΔ monoculture, (C) co-culture *wildtype* (trans well) versus *C. albicans wildtype* monoculture. (D) co-culture *wildtype* versus *S aureus* monoculture (unbuffered), (E) co-culture als1/als3 ΔΔ/ΔΔ versus *S aureus* monoculture (unbuffered), (F) co-culture *wildtype* (trans well) versus *S aureus* monoculture (unbuffered), (G) *S aureus* monoculture (100 mM HEPES) versus *S aureus* monoculture (unbuffered). N-ECVFs are coloured green and non ECVF proteins light grey. Significance lines are based on either a FDR of 0.05 (dashed) or 0.01 (full line). Proteins above the FDR 0.05 line were considered significantly differing.

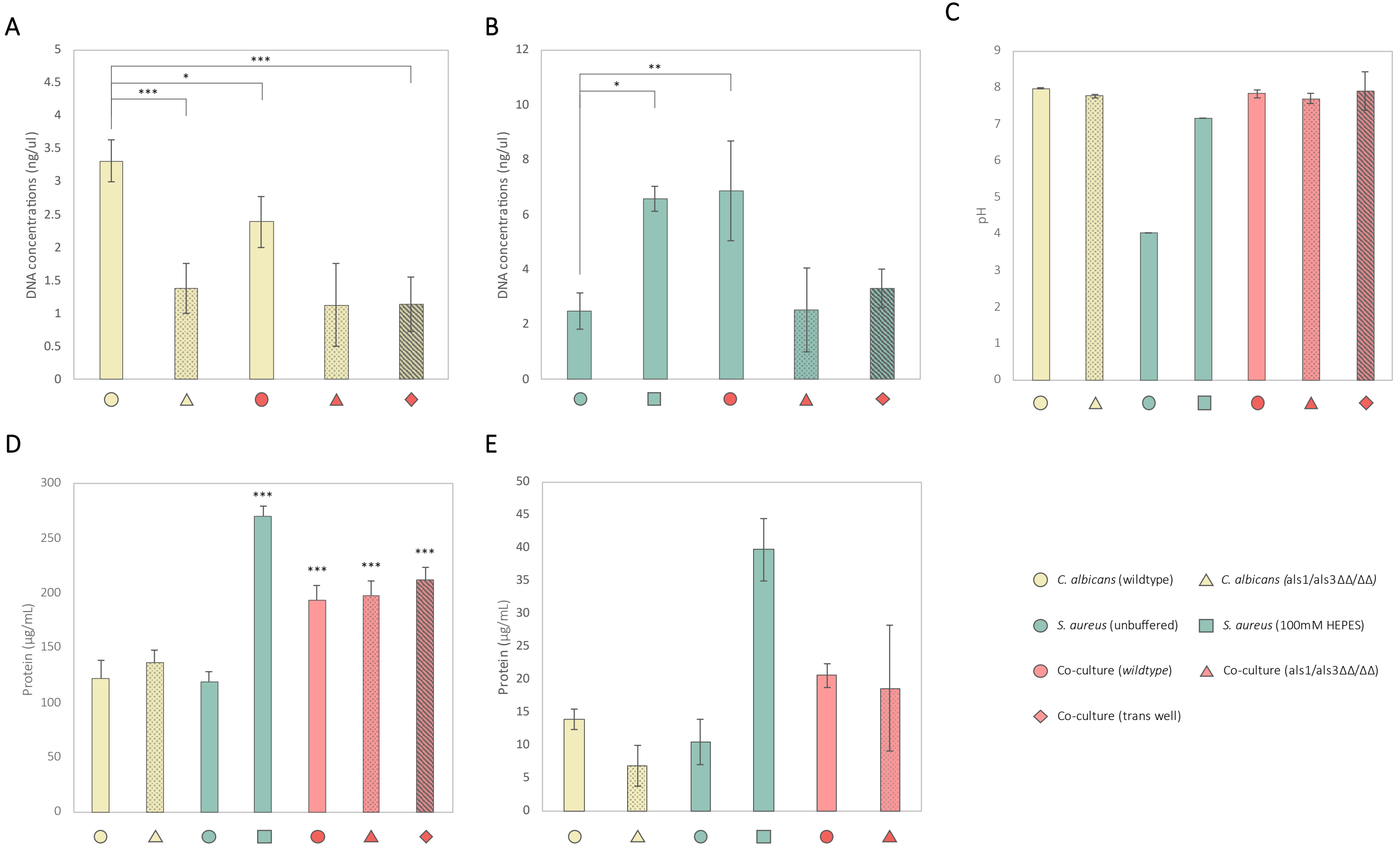


**Figure S3. Basic characterization of the cultures under different conditions.** (A) fungal and (B) bacterial DAN concentrations of the biofilms from which the secretomes were extracted. (C) The pH values of the spent medium as measured before protein concentration. (D) protein concentration of the concentrated secretome as measured before mass spectrometry measurement. (E) protein concentration of non-concentrated secretomes as measured before Ca 9-22 and HO1N1 cytotoxicity experiments.

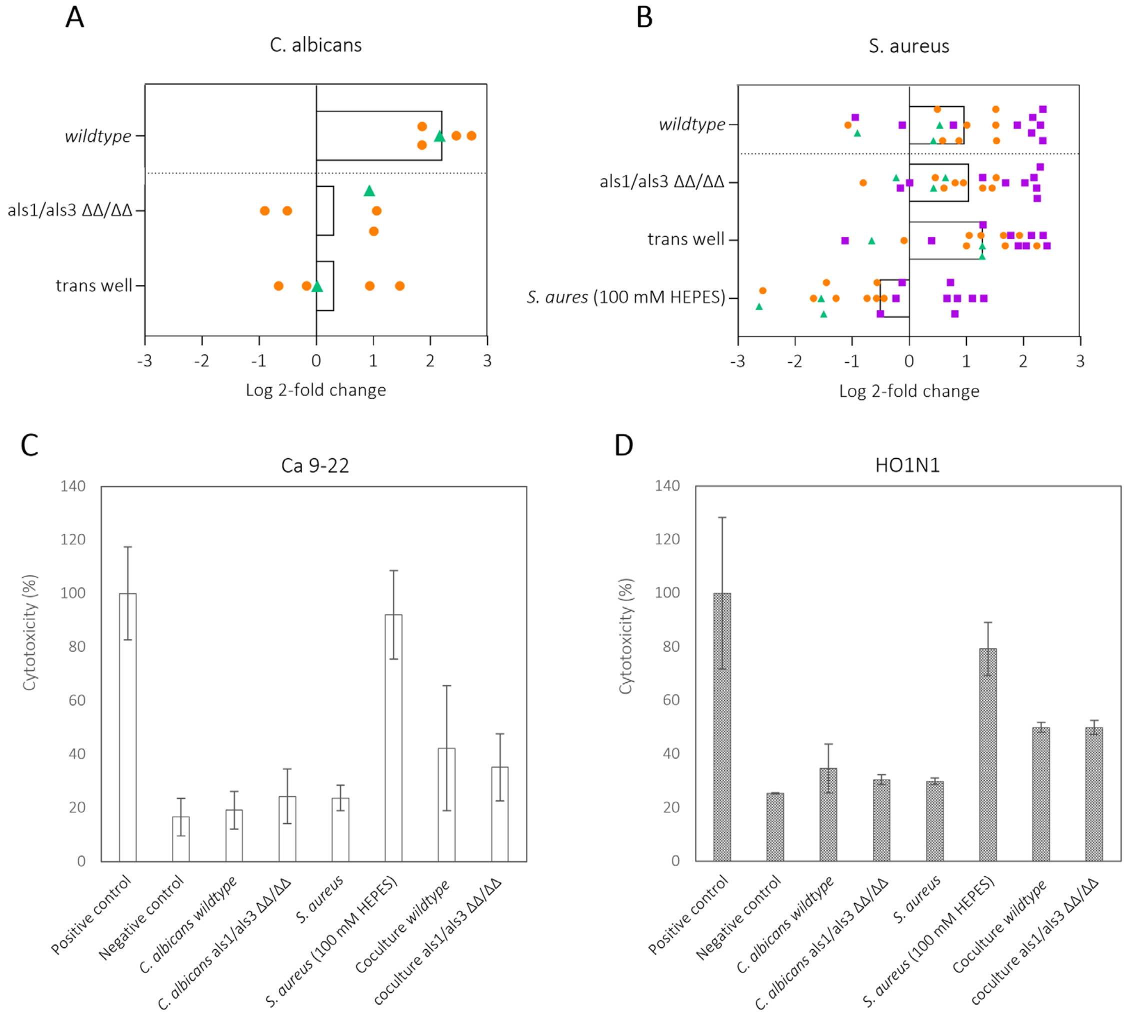


**Figure S4. Cytotoxicity of *C. albicans* and *S. aureus* secretomes increases during co-culturing.** Bar plots of summed log 2-fold changes of damaging ECVFs of (A) *C. albicans* and (B) *S. aureus*, separated by culture condition. Orange circles represent proteolytic ECVFs, green triangles lipolytic ECVFs, and purple squares represent cytolytic ECVFs. Cytotoxicity of *C. albicans* and *S. aureus* mono and co-culture secretomes towards (C) Ca 9-22 (gingival squamous) (n=2) and (D) HO1N1 (buccal mucosa squamous) (n=1) cells. Error bars represent standard deviation from two biological and three technical replicates concerning Ca 9-22 cells and three technical replicates of HO1N1 cells.
